# The impact of allometry on vomer shape and its implications for the taxonomy and cranial kinesis of crown-group birds

**DOI:** 10.1101/2020.07.02.184101

**Authors:** Olivia Plateau, Christian Foth

## Abstract

Crown birds are subdivided into two main groups, Palaeognathae and Neognathae, that can be distinguished, among other means, by the organization of the bones in their pterygoidpalatine complex (PPC). Shape variation of the vomer, which is the most anterior part of the PPC, was recently analysed with help of geometric morphometrics to discover morphological differences between palaeognath and neognath birds. Based on this study, the vomer was identified as sufficient to distinguish the two main groups (and even some inclusive neognath groups) and their cranial kinetic system. As there are notable size differences between the skulls of Palaeognathae and Neognathae, we here investigate the impact of allometry on vomeral shape and its implication for taxonomic classification by re-analysing the data of the previous study. Different types of multivariate statistical analyses reveal that taxonomic identification based on vomeral shape is strongly impaired by allometry, as the error of correct identification is high when shape data is corrected for size. This finding is evidenced by a great overlap between palaeognath and neognath subclades in morphospace. Correct taxonomic identification is further impeded by the convergent presence of a flattened vomeral morphotype in multiple neognath subclades. As the evolution of cranial kinesis has been linked to vomeral shape in the original study, the correlation between shape and size of the vomer across different bird groups found in the present study questions this conclusion. In fact, cranial kinesis in crown birds results from the loss of the jugal-postorbital bar in the temporal region and ectopterygoid in the PPC and the combination of a mobilized quadratezygomatic arch complex and a flexible PPC. Therefore, we can conclude that vomer shape itself is not a suitable proxy for exploring the evolution of cranial kinesis in crown birds and their ancestors. In contrast, the evolution of cranial kinesis needs to be viewed in context of the braincase, quadrate-zygomatic arch and the whole pterygoid-palatine complex.

## Introduction

The pterygoid-palatine complex (PPC) of crown birds is mainly formed by five bones: the unpaired vomer that results from the fusion of the originally paired vomer elements, and the paired pterygoids and palatines. The general morphology of the PPC was first studied by **Huxley (1867)**, who distinguished the clade Palaeognathae from all other birds on the basis of palatal morphology. Although the PPC of Palaeognathae is quite variable (**McDowell, 1948**), it is characterized by a large vomer that is only partly fused. The pterygoids and palatines are highly connected, forming a rigid unit that articulates with the braincase via well-developed basipterygoid processes, while a contact with the parasphenoid is not present (see **Bellairs and Jenkin, 1960**; **Zusi, 1993**; **Gussekloo et al**., **2001**; **Mayr, 2017**; **Fig. 1A**). In contrast, neognath birds possess a movable joint between pterygoid and palatine, which plays an important role in the kinematic movement of the upper jaw. Here, the pterygoid articulates with the parasphenoid, while the basipterygoid processes are often reduced. The vomer is highly variable in size and shape and often has no connection with the upper jaw beyond an association with the nasal septum and the palatine. In some neognaths, the vomer is greatly reduced or even absent (see **Bellairs and Jenkin, 1960**; **Bock, 1964**; **Zusi, 1993**; **Mayr, 2017**; **Fig. 1A**).

**Figure 1.**
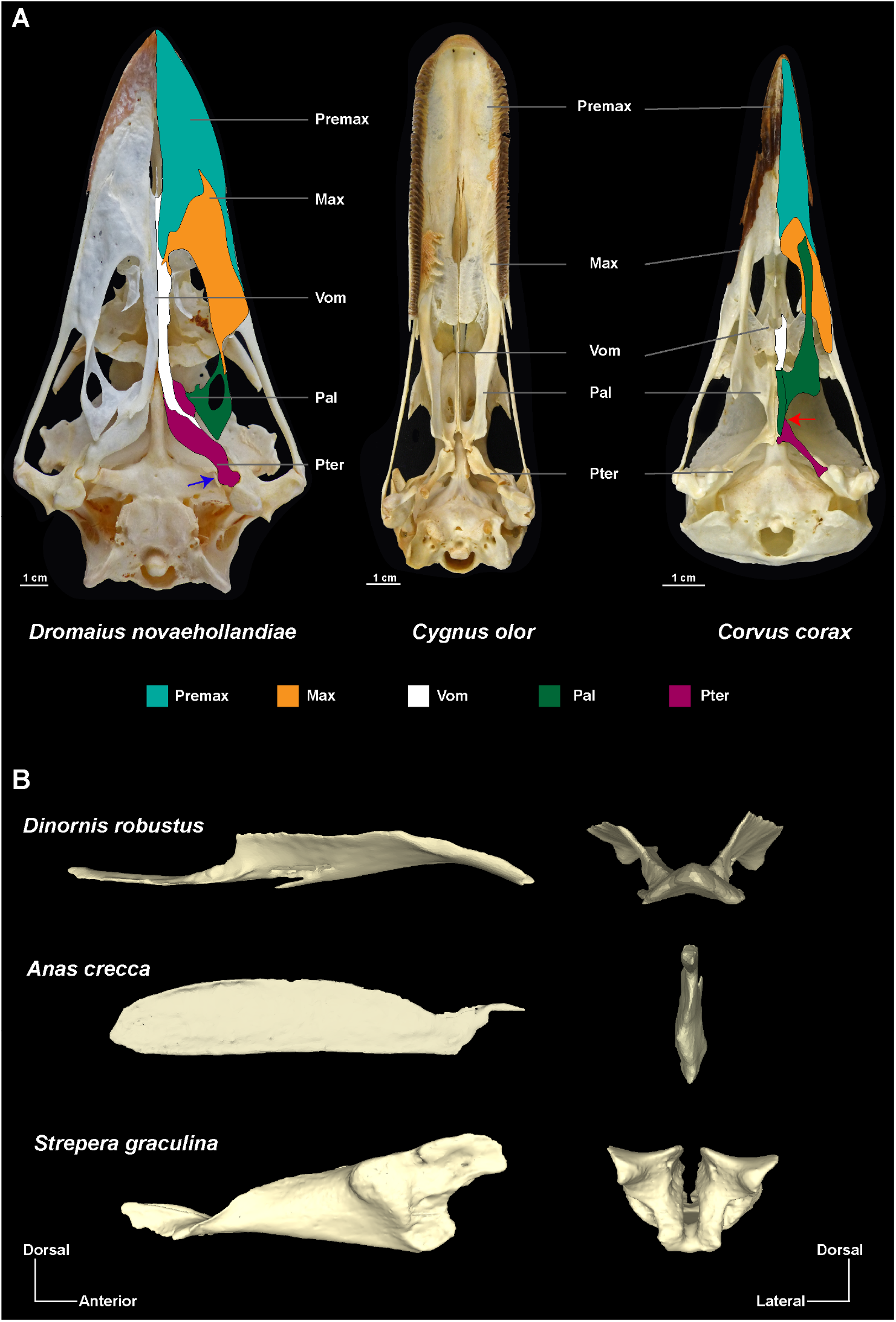
Anatomical organization of the pterygoid-palatine complex (PPC) and shape variability of the vomer in palaeognath and neognath birds. (A) Palates of *Dromaius novaehollandia* (left), *Cygnus olor* (middle) and *Corvus corax* (right) in ventral view (all specimens from the Natural History Museum of Fribourg/University of Fribourg). For *Dromaius novaehollandiae* and *Corvus corax* the main organization of palate morphology is highlighted in a coloured scheme. The blue arrow in *Dromaius novaehollandiae* indicates the contact between the basipterygoid process and the pterygoid. The red arrow in *Corvus corax* indicates the mobile joint between pterygoid and palatine. (B) 3D models of the vomer of *Dinornis robustus, Anas crecca* and *Strepera graculina* in lateral view (left) and anterior view (right), not at scale (3D models from **Hu et al**., **2019**). Abbreviations: Max, Maxillary; Pal, Palatine; Premax, Premaxilary; Pter, Pterygoid; Vom, Vomer.

In a recent paper, **Hu et al. (2019)** investigated palate evolution in crown birds and their stem, focusing on the morphology of the vomer. Using 3D geometric morphometrics, the study found that the vomeral shape of neognaths is clearly distinguishable from Palaeognathae, in that the latter group has a stronger similarity with their non-avian ancestors. Linking vomer shape with the kinetic abilities of the skull, the authors concluded that cranial kinesis represents an innovation of Neognathae. Furthermore, the authors concluded that vomeral morphology allows for a taxonomic differentiation between the major groups of Neognathae, namely Aequorlitornithes, Galloanserae, Gruiformes, and Inopinaves. However, according to their *PCA* results, all groups strongly overlap each other within PC1, while a taxonomic differentiation is only noticeable within PC2 (other principal components are not shown). Taking the great size variation of the vomer of neognath birds into account (**Zusi, 1993**), we wonder if the reported taxonomic differentiation between palaeognaths and the neognath subclades could alternatively be related to allometry, i.e. the dependence of shape on size (**Klingenberg, 1998**), rather than pure shape variation. In order to test this hypothesis, we re-analysed the dataset of **Hu et al. (2019)**, comparing allometric shape data with non-allometric residuals, and re-evaluating the role of the vomer in the evolution of cranial kinesis in crown birds.

## Material and methods

The published 3D models and landmarks data of 41 specimens including 36 species were downloaded from **Hu et al. (2019)** (10.6084/m9.figshare.7769279.v2). This dataset contains five extinct species (two stem line representatives: the troodontid *Sinovenator changii*, the Avialae *Sapeornis chaoyangensis*; and three fossil palaeognath crown birds from the clade Dinornithiformes: *Pachyornis australis, Megalapteryx didinus* and *Diornis robustus*), five extant Paleognathae and 27 extant Neognathae representing the two major clades of crown birds.

The original data (Dataset A) is composed of five anatomical landmarks and 43 semi-landmarks (see **Hu et al**., **2019**). The landmark data were imported into the software R v.3.5.2 (R **Core Team, 2011**). Using the *plotAllSpecimens* function of Geomorph v.3.2.1 (**Adams and Otárola-Castillo, 2013**) in R, we notice great variability for each anatomical landmark, resulting from two main shapes of the vomer. The majority of bird possesses a fused vomer that is bilaterally symmetric and roof-shaped in transection, with a horizontal orientation within the pterygoid-palatine complex (**Fig. 1B**). In contrast, some members of Aequorlitornithes (e.g., *Podiceps nigricollis*, and *Podilymbus podiceps*), Galloanserae (e.g., *Anas crecca, Anseranas semipalmata*, and *Cairina moschata*) and Inopinaves (e.g., *Aquila audax, Falco cenchroides*, and *Haliastur sphenurus*) possess a fused vomer that is completely mediolaterally flattened in transection and vertically orientated within the pterygoid-palatine complex (**Fig. 1B**). Therefore, we created a second dataset (Dataset B), where species with this latter, flat vomer morphology were excluded. Furthermore, the palaeognath birds *Struthio camelus* and *Dromaieus novaehollandiae* of the original Dataset A were represented by both juvenile and adult specimens. Because ontogenetic variation could, however, potentially affect size and position of the palaeognath morphospace, we removed the juvenile and subadult specimens of *Struthio camelus* and *Dromaieus novaehollandiae* in order to reran the analysis just with adult semaphoronts (Dataset C). Finally, we created a fourth dataset, where both juvenile/subadult specimens and species with flat vomers were removed from the sample (Dataset D).

For superposition of the 3D landmark data, we followed **Hu et al. (2019)** by performing a *Generalized Procrustes Analysis* (*GPA*). The *GPA* was done with the help of the *gpagen* function in Geomorph. Afterward, we performed a *principal component analysis* (*PCA*) in order to visualize the shape variability of the vomer and the variance of morphospace for two groupings: (1) Paleognathae versus Neognathae and (2) Paleognathae, Inopinaves, Galloanserae, Gruiformes and Aequorlitornithes. This was done with the *plotTangentSpace* function from Geomorph.

Because the vomer showed great variation in centroid size after superimposition, ranging from 14.60 (*Manorina melanocephala*) to 168.32 (*Dromaieus novehollandia*), we tested if there is a significant correlation between Procrustes coordinates and log-transformed centroid size (**Goodall, 1991**) using the function *procD*.*lm* in Geomorph. This function performs a multivariate regression between the shape and size with a permutation of 10,000 iterations. A significant relationship between both parameters indicates that the superimposed shape still contains an allometric signal. Based on this correlation we estimated non-allometric residuals of the Procrustes coordinates and repeated the *PCA*. In addition, we tested each of the first eleven PCs that together describe more than 95 of total variation for allometric signals.

A set of 1,000 relaxed-clock molecular trees, which follow the topology of **Hackett et al. (2008)** and summarize the range of uncertainties in terms of time calibration of ancestral nodes, were downloaded from birdtree (birdtree.org) (**Jetz et al**., **2012**,**2014**) including all extant bird species in the dataset (**Supplementary Data S1**). Due to uncertainties in the taxonomic identification of *Aquila* sp., this specimen was removed from the sample as we could not include it in the phylogeny. Because the specimen occupies almost the same position as *Aquilla audax*, we consider this deletion to have a negligible effect on the outcome of the analyses. Furthermore, the species *Sterna bergii* and *Grus rubincunda* used in the analysis from **Hu et al. (2019)** are junior synonyms of *Thalasseus bergii* (**Bridge et al**., **2005**) and *Antigone rubicunda* (**Krajewski et al**., **2010**). Using the function *consensus*.*edges* in the R package phytools v.0.7-20, we computed a temporal consensus. The extinct dinornithiform species were placed as sister-group to Tinamidae following **Mitchell et al. (2014)**. Because of their recent extinction (**Holdaway and Jacomb, 2000**; **Turvey and Holdaway, 2005**), the age was set to zero, similar to the other crown birds. The stem line representatives *Sinovenator changii* and *Sapeornis chaoyangensis* were added following the time-calibrated phylogeny of **Rauhut et al. (2019)**. Because of the presence of juvenile specimens in datasets A and B, we added the juvenile specimens by splitting OTU (operational taxonomic unit) of *Struthio camelus* and *Dromaieus novehollandia* into a polytomy with each sub-OTU having a branch length of one year (this value had to be standardized, as *pFDA* requires an isometric tree). The impact of phylogeny on shape and centroid size (log-transformed) was tested using the function *physignal* in Geomorph. Based on *K*-statistics, this method evaluates the impact of phylogeny on a multivariate dataset relative to what is expected, if evolution is simulated under a Brownian motion model (**Blomberg et al**., **2003**).

To explore the size of the single morphospaces of each taxon, we computed the morphological disparity using *morphol*.*disparity* in Geomorph. This analysis uses the Procrustes variance as disparity metric, which is the sum of all diagonal elements of a group covariance matrix divided by the number of observations within the group (**Zelditch et al**., **2012**). Statistical comparisons of group sizes were executed with help of a permutation test with 999 iterations. To test for potential overlap in morphospace of vomer shapes in different clades of crown bird (see grouping 1 and 2) and their relation to the stem line representatives *Sinovenator changii* and *Sapeornis chaoyangensis*, we applied three different multivariate statistical methods, using the first eleven PCs as input data. We first applied a nonparametric multi-variate analysis of variance (*perMANOVA*). This method evaluates the potential overlapping of groups in morphospace by testing the significance of their distribution on the basis of permutation (10,000 replications) and Euclidean distance (as one of several possible distance measures), not requiring normal distribution of the data (**Anderson, 2001**; **Hammer and Harper, 2006**). The spatial relationship of groups relative to each other is expressed by an *F* value and *p* value. For the five-group comparison, the *p* values were Bonferroni-corrected by multiplying the value with the number of comparisons. Next, we ran a discriminant analysis (*DA*), which reduces a multivariate data set to a smaller set of dimensions by maximizing the separation between two or more groups using Mahalanobis distances. This distance measure is estimated from the pooled within-group covariance matrix, resulting in a linear discriminant classifier and an estimated group assignment for each species. The results were cross-validated using Jackknife resampling (**Hammer and Harper, 2006**; **Hammer, 2020**). Both multivariate tests were done with the program PAST v.4.03 (**Hammer et al**., **2001**). Finally, we performed a phylogenetic flexible discriminant analysis (*pFDA*) (**Schmitz and Motani, 2011**; **Motani and Schmitz, 2011**) in R. This method removes the phylogenetic bias from the categorical variables before the actual discriminant analysis is undertaken by estimating Pagel’s lambda, which tests how the grouping correlates with phylogeny. This was done for all allometric and non-allometric datasets.

The error of correct identification from the resulting confusion matrices was compared between allometric and non-allometric data. For these comparisons, we used nonparametric *Mann-Whitney U* and *Kruskal-Wallis* tests, which both estimates, whether or not two univariate samples were taken from populations with equal medians, being more robust against small sample sizes and data without normal distribution than parametric tests (**Hammer and Harper, 2006**). Both tests were carried out with PAST.

Finally, we applied an ordinary least square regression analysis to 19 species, testing the correlation between log-transformed vomer size and the skull size using a log-transformed box volume (Height x Width x Length). The measurements of the skull box volume were taken from skullsite (skullsite.com).

## Results

Based on the *PCA* of the original dataset, the first two PCs explain over 52% (**Fig. 2A**) of total shape variation (PC1: 27.5%; PC2: 25.1%). The morphospace of Palaeognathae and Neognathae is almost equal in size. Taking the small sample size of Palaeognathae into account, the size of their morphospace indicates relatively great shape variation. This is supported by the Procrustes variances, which indicates a larger shape disparity in Palaeognathae. Both Palaeognathae and Neognathae show a strong overlap along PC1 and a partial overlap along PC2. When comparing neognath subclades, Aequorlitornithes show strong overlap along both PCs with the palaeognath morphospace. Gruiformes lie in the overlapping area of both groups. The morphospace of Inopinaves and Galloanserae overlap with each other on both axes, but are separated from Palaeognathae, Aequorlitornithes and Gruiformes along PC2. Within Neognathae, Galloanserae have the largest shape disparity, becoming successively smaller in Inopinaves, Aequorlitornithes and Gruiformes (**Supplementary Data S2; S3: Fig. S1; S4: Table S1**).

**Figure 2.**
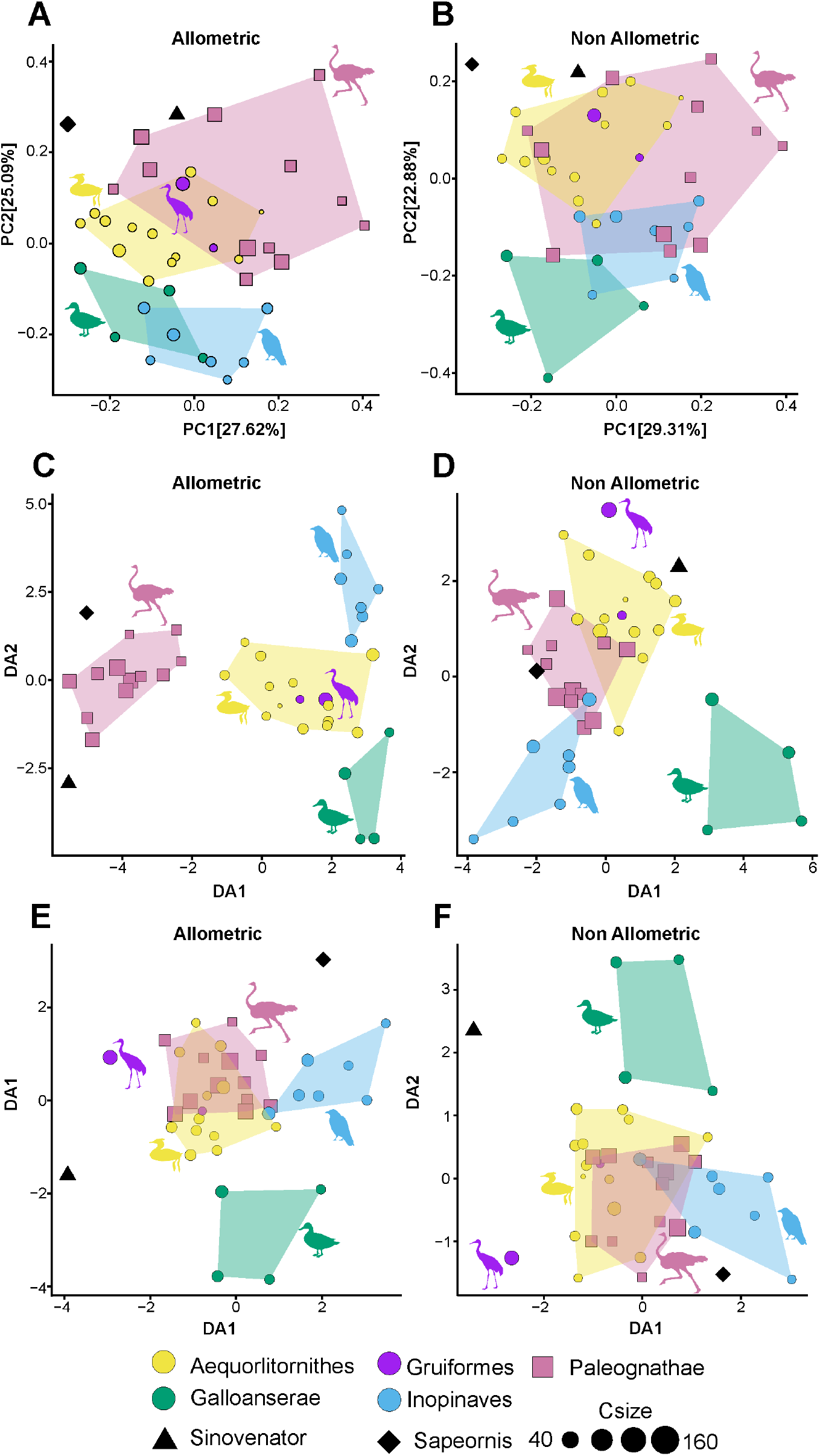
Differences between allometric and non-allometric morphospaces of the vomer (Dataset A) in palaeognath and neognath birds. (A) *PCA* (*Principal Component Analysis*) results of allometric data. (B) *PCA* results of non-allometric data. (C) *DA* (*Discriminant Analysis*) results of allometric data. (D) *DA* results of non-allometric data. (E) *pFDA* (*phylogenetic Flexible Discriminant Analysis*) results of allometric data. (F) *pFDA* results of non-allometric data. Centroid sizes (Csize) are indicated by the size of the symbols. All silhouettes are taken from www.phylopic.org/.

Allometry summarizes about 6.4% of total shape variation. Using non-allometric residuals PC1 explains 29.3% and PC2 22.9% (**Fig. 2B**). While the general distribution of the single bird clades does not change along PC1, the groups are less separated along PC2, which contains the major allometric signal within the principal components (slope: -0.523; *R*^2^: 0.185; *p*: 0.005; predicted variation: 19.5%), which is 4.9% of total shape variation in the original dataset. Here, the palaeognath morphospace overlaps fully with Aequorlitornithes and Gruiformes, partly with Inopinaves and marginally with Galloanserae. For the three other datasets, we observe more or less similar general trends before and after size correction, although the single morphospaces are partly shrunk. In all cases, the two stem line representatives *Sinovenator changii* and *Sapeornis chaoyangensis* lie within the marginal area of the palaeognaths/aequorlitornithines morphospace. Here, vomer morphology of the troodontid *Sinovenator changii* is more bird-like than that of the avialian *Sapeornis chaoyangensis*. The exclusion of the allometric shape variation has only a minor impact on disparity. Thus, all disparity trends found in the original dataset persist in the non-allometric datasets (**Supplementary Data S2; S3: Fig. S1; S4: Table S1**).

As previously detected by **Hu et al. (2019)**, vomer shape is impacted by phylogeny. Neither the modification of the sampling nor the exclusion of allometry changes this relationship. In contrast, log centroid size does not contain a phylogenetic signal (**Supplementary Data S4: Table S2**).

In all studied datasets, the *perMANOVA* found a significant separation between palaeognath and neognath birds, showing no impact of allometry. For the five-group comparison of the original dataset (A), the overall results still indicate significant separation between clades for both allometric and non-allometric data. However, group-by-group comparison of allometric data indicates an overlap in morphospace of Gruiformes with Aequorlitornithes, Inopinaves, Galloanserae and Palaeognathae. These overlaps of Gruiformes with other bird clades persist when allometry is removed from shape, but in addition, Aequorlitornithes cannot be distinguished from Palaeognathae anymore, as indicated by the *PCA* results (**Fig. 2A, B**). The overlap between clades increases with the exclusion of species with flat vomers and non-adult semaphoronts (**Supplementary Data S4: Tables S3, S4**).

For the original dataset (A) with allometry included, the *DA* identifies all species correctly as Palaeognathae or Neognathae. The error of false identification increases to 2.6% if the data are jackknifed. When allometry is removed, the error increases to 13.2% before and 36.8% after jackknife resampling. In the former case, the misidentifications are restricted to neognath birds, which are wrongly classified as Palaeognathae, while jackknifing leads to identification errors in both groups. For the five-group comparison, all species of dataset (A) are correctly identified, when allometry is still present. The error is 18.4% after jackknife resampling, showing minor mismatches in all clades. Excluding allometry, the error increases to 10.5% before and 47.4% after jackknifing. While in the former case, a few Aequorlitornithes (2) and Inopinaves (1) species are wrongly identified as Palaeognathae (**Fig. 2C, D**), Palaeognathae cannot be separated from the neognath subclades anymore after resampling. The exclusion of species with flat vomers and non-adult semaphoronts leads to an increase of error (**Supplementary Data S4: Tables S3–S6**).

The *pFDA* found 15.8% of incorrect identifications when Palaeognathae are compared with neognaths in the original dataset (A). This error increases to 31.6% if shape is corrected for allometry. In both cases, error is primarily based on the incorrect identification of palaeognath specimens as neognaths. When Palaeognathae are compared with neognath subclades, the error of correct identification is 10.5% before and 26.3% after allometry is removed from the data. For the allometric data, the misidentifications result from the overlap between Paleognathae, Aequorlitornithes and Gruiformes. The misidentifications between these three groups are increased when shape is corrected for allometry, while Inopinaves are in part also wrongly identified as Palaeognathae. The exclusion of species with a flat vomer and/or non-adult semaphoronts usually causes a decrease of false identifications. However, the non-allometric dataset shows an increase in error for the two-group comparison, when species with flat vomers are excluded, and for the five-group comparison, when only adult semaphoronts are taken into account (**Fig. 2E, F**). Nevertheless, for all four datasets, the error of correct identification is significantly higher for non-allometric vomer shapes (**Fig. 3A; Supplementary Data S4: Tables S3–S6**).

**Figure 3.**
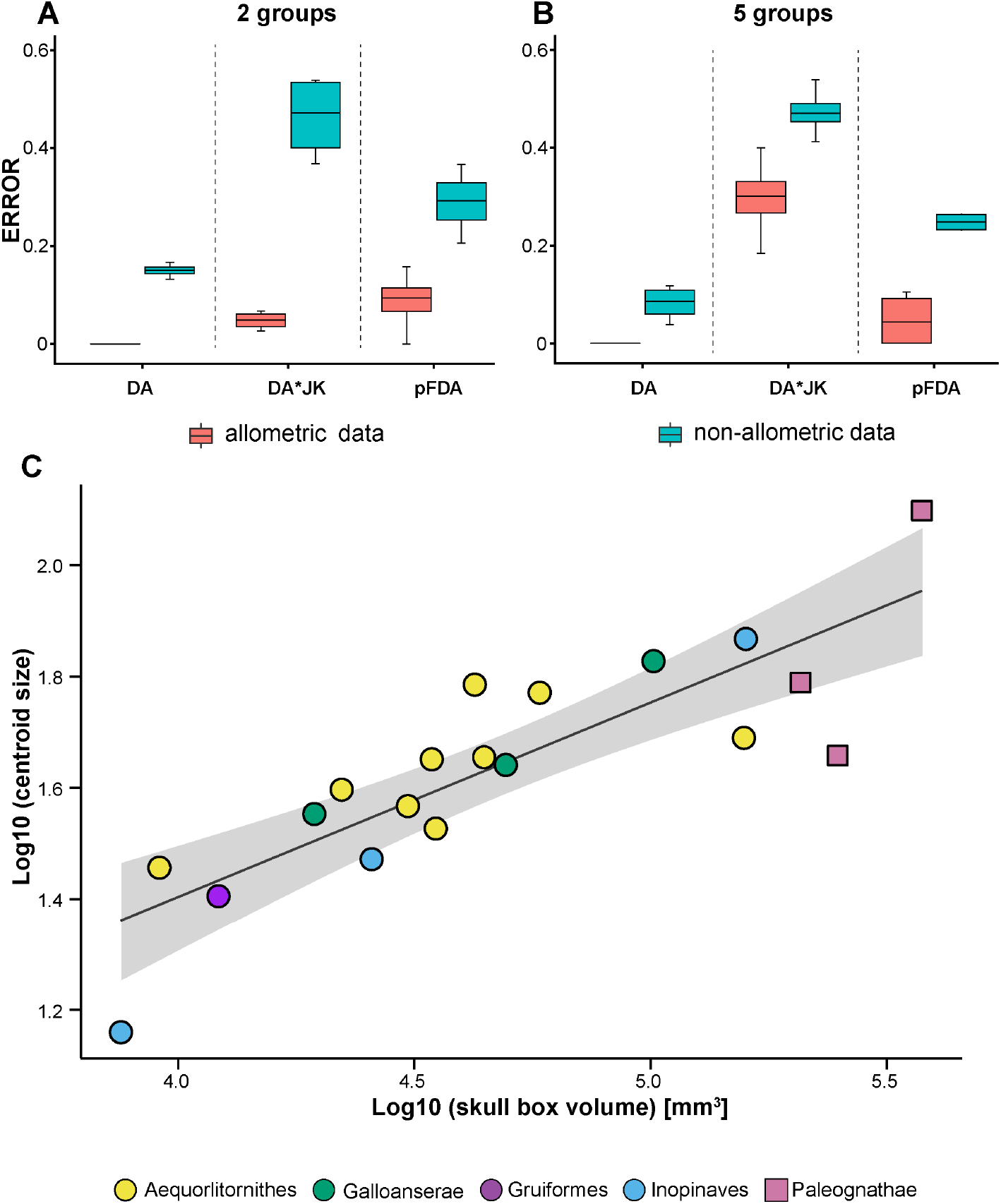
Errors of correct taxonomic identification for all comparisons of Datasets A–D. Two-group identification (Palaeognathae and Neognathae) before (red) and after (green) correction for allometry. *DA, Discriminant analysis*; *DA*JK, Discriminant Analysis with jackknife resampling*; *pFDA, phylogenetic Flexible Discriminant Analysis*. (B) Five-group identification (Palaeognathae, Aequorlitornithes, Galloanserae, Gruiformes and Inopinaves). (C) *OLS* regression (black line) between log-transformed skull box volume and logtransformed centroid size of the vomer. Grey shadow marks the area of the 95% confidence interval.

## Discussion

The skull of crown birds possesses a complex kinetic system that includes a mobilized quadrate, the zygomatic arch (= jugal bar) and the pterygoid-palatine complex (PPC) that allows for the simultaneous, but restricted motion of both jaws (**Bock, 1964**; **Zusi, 1984**). According to **Zusi (1984)**, the kinetic system can be differentiated into three main types: (1) prokinesis describes the rotation of the whole beak around the nasal-frontal hinge; (2) amphikinesis is derived from prokinesis, including the rotation of the beak around the nasalfrontal hinge plus an additional flexion of the anterior portion of the beak; and (3) rhynchokinesis, which in contrast includes a simple flexion of the beak around one or several bending zones rostral to the nasal-frontal suture, lacking a true hinge. Depending on the position of the bending zones, rhynchokinesis can be further differentiated into five subtypes (**Zusi, 1984**). Most palaeognath birds possess central rhynchokinesis, while neognaths have realized all types of cranial kinesis (**Zusi, 1984**), including some taxa with akinetic skulls (**Reid, 1835**; **Sims, 1955**; **Degrange et al**., **2010**). In the past, several authors (**Hofer, 1954**; **Simonetta, 1960**; **Bock, 1963**) suggested a close relationship between the morphology of the PPC and type of cranial kinesis. However, **Gussekloo et al. (2001)** demonstrated that all types of kinesis present in crown birds have similar movements of the quadrate, zygomatic arch and PPC. Palaeognathae and Neognathae only differ in the magnitude of kinetic movements in that Palaeognathae have slightly more restricted movement due to their rigid palate missing a movable joint between the pterygoid and palatine (**Gussekloo and Bout, 2005**).

Thus, although the results of geometric morphometric analysis of the vomer shape by **Hu et al. (2019)** imply at first glance a distinct separation between Palaeognathae and Neognathae, this separation does not necessarily reflect their conclusions regarding the evolution of cranial kinesis in crown birds, i.e., that cranial kinesis represents an innovation of Neognathae. As indicated by the *PCA*, Palaeognathae occupy an enormous vomeral morphospace (**Hu et al**., **2019**), which mirrors their generally large palatal disparity (see **McDowell, 1948**) and partly overlaps with Gruiformes and Aequorlitornithes. In all cases tested, however, the exclusion of allometric shape variation generally increases the error of misidentification between all groups (**Fig. 3A; Supplementary Data S4: Table S7**), indicating that the taxonomic distinctions of shape found by **Hu et al. (2019)** are at least partly an artefact of size. This primarily concerns PC2, which according to **Hu et al. (2019)** separates Palaeognathae from Neognathae, but also contains the major part of allometric information. According to shape variation explained by PC2, larger birds tend to evolve vomers that are more dorsoventrally compressed. Only members of the Galloanserae could still be identified with a high amount of certainty when allometry is excluded.

Thus, our finding supports previous studies that demonstrated a relevant impact of allometry on skull shape evolution in birds (**Klingenberg and Marugán-Lobón, 2013**; **Bright et al**., **2016**; **Linde-Medina, 2016**; **Tokita et al**., **2017**; **Bright et al**., **2019**). By modifying the dataset, it becomes further clear that both the homoplastic presence of flat vomers in Aequorlitornithes, Inopinaves, Galloanserae (Dataset B) and ontogenetic variation (Dataset C) affects the accuracy of taxonomic identification. In addition, Palaeognathae and Neognathae do not differ in vomer size when compared to the head size (**Fig. 3B**). Consequently, vomer shape is not practical for taxonomic identification and should not be used as a proxy to infer the presence or absence of cranial kinesis in crown birds or their stem. As the manifold shape diversity of crown bird’s skulls is impaired by a tessellated evolution with multiple convergent events (e.g., **Zusi, 1993**; **Felice and Goswami, 2018**), the use of isolated elements for taxonomic identification and/or biomechanical implications should be treated generally with some caution.

In fact, *DA* and *pFDA* frequently identified the troodontid *Sinovenator changii* and avialan *Sapeornis chaoyangensis* as neognaths or neognath subclades when allometry is excluded, while the original dataset implied a referral to Palaeognathae (see also **Hu et al**., **2019**). However, the skull anatomy of both species indicates no cranial kinesis (**Xu et al**., **2002**; **Wang et al**., **2017**; **Yin et al**., **2018**; **Hu et al**., **2020**).

The origin and evolution of cranial kinesis in the stem line of birds is still not well understood due to the rarity of complete three-dimensional skulls. However, skull material from the ornithurines *Ichthyornis dispar* and *Hesperornis regalis* indicates a certain degree of rhynchokinesis (**Bühler et al**., **1988**; **Field et al**., **2018**) that might be comparable to that of extant Palaeognathae or some Aequorlitornithes, but further shows that this functional character was already present before the origin of the crown. Their kinesis is indicated by the loss of the jugal-postorbital bar and the ectopterygoid (resulting in a loss of contact in the jugal with the skull roof and the palate), the presence of a mobile bicondylar quadrate and a mobile joint between quadrate and quadratojugal. Recently, **Plateau and Foth (2020)** speculated that the peramorphic bone fusion in the braincase could be also related to cranial kinesis, in which the fusion-induced immobility constrains a controlled kinetic dorsoventral flexion of the avian beak during biting/picking. Based on these characters, most Mesozoic Avialae (including *Sapeornis chaoyangensis*) still had akinetic skulls, although some Enantiornithes possessing a reduced jugal-postorbital bar might have evolved primitive kinesis convergently to Ornithurae (**O’Connor and Chiappe, 2011**).

In summary, allometry describes the relationship between size and shape, which is one of the key concepts in biology to explain variation of shape in organisms (**Klingenberg, 1998**), and is crucial for studying taxonomy, ontogeny and functional morphology. Investigating the effect of allometry on the vomer shape in crown birds with help of multiple multivariate statistical methods, indicates that this bone is not a good proxy for taxonomy. Shape differences between Palaeognathae and Neoganthae are clearly affected by size and can neither be used to differentiate between different types of cranial kinesis nor to explain the evolution of cranial kinesis within crown birds. In contrast, the evolution of cranial kinesis in birds needs to be studied in context of the whole pterygoid-palatine complex and its contacts with the braincase and quadrate-zygomatic arch.

## Acknowledgements

We thank Walter Joyce, Roland Sookias and Sergio Martínez Nebreda for their critical comments on previous versions of the manuscript. The Swiss National Science Foundation is thanked for its financial support (PZ00P2_174040 to C.F.).

## Additional information

### Funding

This study was funded by the Swiss National Science Foundation (PZ00P2_174040).

### Competing interests

The authors declare they have no personal or financial conflict of interest relating to the content of this study.

## Author contributions

O.P. and C.F. designed the research project and analysed the data; O.P. and C.F. wrote the paper and prepared all figures. All authors have seen and approved the manuscript.

## Data availability

The 3D models and landmarks data of **Hu et al. (2019)** are available at Figshare (DOI: 10.6084/m9.figshare.7769279.v2).

The data relevant to the present study are provided below as **Supplementary information**.

## Supplementary information

All files are available online (doi: 10.1101/2020.07.02.184101).

1. Data S1: Phylogenetic trees used for *pFDA*.
2. Data S2: *PCA* results of all datasets before and after correction for allometry.
3. Data S3: *PCA* plots of Datasets B-D before and after correction for allometry.
4. Data S4: Results of *npMANOVA, DA* and *pFDA*.
5. Data S5: R Code including all statistical analyses.

## Notes

https://doi.org/10.6084/m9.figshare.7769279.v2

### Competing Interest Statement

The authors have declared no competing interest.

