## Supplementary Data S3 for "The impact of allometry on vomer shape and its implications for the taxonomy and cranial kinesis of crown-group birds"

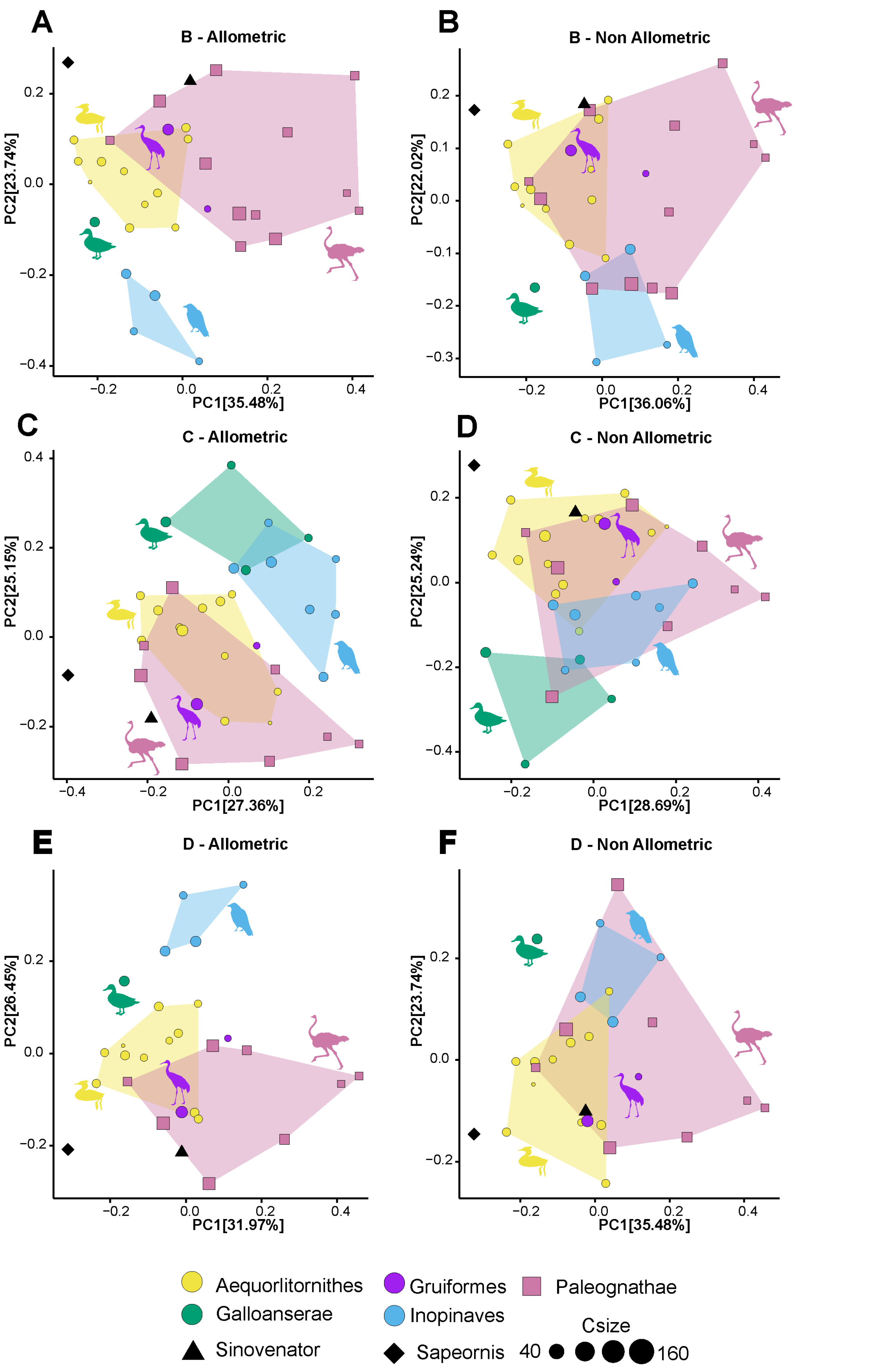
A
B - Allometric
PC2[23.74%]
PC1[35.48%]
B
B - Non Allometric
PC2[22.02%]
PC1[36.06%]
C
C - Allometric
PC2[25.15%]
PC1[27.36%]
D
C - Non Allometric
PC2[25.24%]
PC1[28.69%]
E
D - Allometric
PC2[26.45%]
PC1[31.97%]
F
D - Non Allometric
PC2[23.74%]
PC1[35.48%]
Aequorlitornithes
Galloanserae
Sinovenator
Gruiformes
Inopinaves
Sapeornis
Paleognathae
Csize
40
160

**Supplementary Fig. S1. Differences between allometric and non-allometric morphospaces of the vomer in palaeognath and neognath birds. (A)** *PCA* (*Principal component analysis*) results of (Dataset B) with allometric data. **(B)** *PCA* results of (Dataset B) with non-allometric data. **(C)** *PCA* results of (Dataset C) with allometric data. **(D)** *PCA* results of (Dataset C) with non-allometric data. **(E)** *PCA* results of (Dataset D) with allometric data. **(F)** *PCA* results of (Dataset D) with non-allometric data. Centroid sizes (Csize) are indicated by the size of the symbols. All silhouettes are taken from http://www.phylopic.org/.
